# Shifting spaces: which disparity or dissimilarity metrics best summarise occupancy in multidimensional spaces?

**DOI:** 10.1101/801571

**Authors:** Thomas Guillerme, Mark N. Puttick, Ariel E. Marcy, Vera Weisbecker

## Abstract

1. Multidimensional analysis of traits are now a common toolkit in ecology and evolution and are based on trait-spaces in which each dimension summarise the observed trait combination (a morphospace or an ecospace). Observations of interest will typically occupy a subset of this trait-space, and researchers will apply one or more metrics to quantify the way in which organisms “inhabit” that trait-space. In macroevolution and ecology these metrics are referred to as disparity or dissimilarity metrics and can be generalised as space occupancy metrics. Researchers use these metrics to investigate how space occupancy changes through time, in relation to other groups of organisms, and in response to global environmental changes, such as global warming events or mass extinctions. However, the mathematical and biological meaning of most space occupancy metrics is vague with the majority of widely-used metrics lacking formal description.
2. Here we propose a broad classification of space occupancy metrics into three categories that capture changes in volume, density, or position. We analyse the behaviour of 25 metrics to study changes in trait-space volume, density and position on a series of simulated and empirical datasets.
3. We find no one metric describes all of trait-space but that some metrics are better at capturing certain aspects compared to other approaches and that their performance depends on both the trait-space and the hypothesis analysed. However, our results confirm the three broad categories (volume, density and position) and allow to relate changes in any of these categories to biological phenomena.
4. Since the choice of space occupancy metric should be specific to the data and question at had, we introduced `moms`, a user-friendly tool based on a graphical interface that allows users to both visualise and measure changes space occupancy for any metric in simulated or imported trait-spaces. Users are also provided with tools to transform their data in space (e.g. contraction, displacement, etc.). This tool is designed to help researchers choose the right space occupancy metrics, given the properties of their trait-space and their biological question.

## Introduction

Groups of species and environments share specific, easily recognisable, correlated characteristics of many kinds: guilds or biomes with shared phenotypic, physiological, phylogenetic or behavioural traits. Organisms or environments should therefore be studied as a set of traits rather than some specific traits in isolation (Donohue et al. 2013; Hopkins and Gerber 2017). Biologists have increasingly been using ordination techniques (see Legendre and Legendre 2012 for a summary) to create multidimensional trait-spaces to either explore properties of the data or test hypotheses (Oksanen et al. 2007; Adams and Otárola-Castillo 2013; Bonhomme et al. 2014; Blonder 2018; Guillerme 2018). For example, in palaeobiology, Wright (2017) use trait-spaces to study how groups of species’ characteristics change through time; in ecology, Jones et al. (2015) study evidence of competition by looking at trait overlap between two populations. However, different fields use a different set of terms for such approaches (Table 1). Nonetheless, they are the same mathematical objects: matrices with columns representing an original or transformed trait value and rows representing observations, such as taxon, field site, etc. (Guillerme 2018).

**Table 1:**
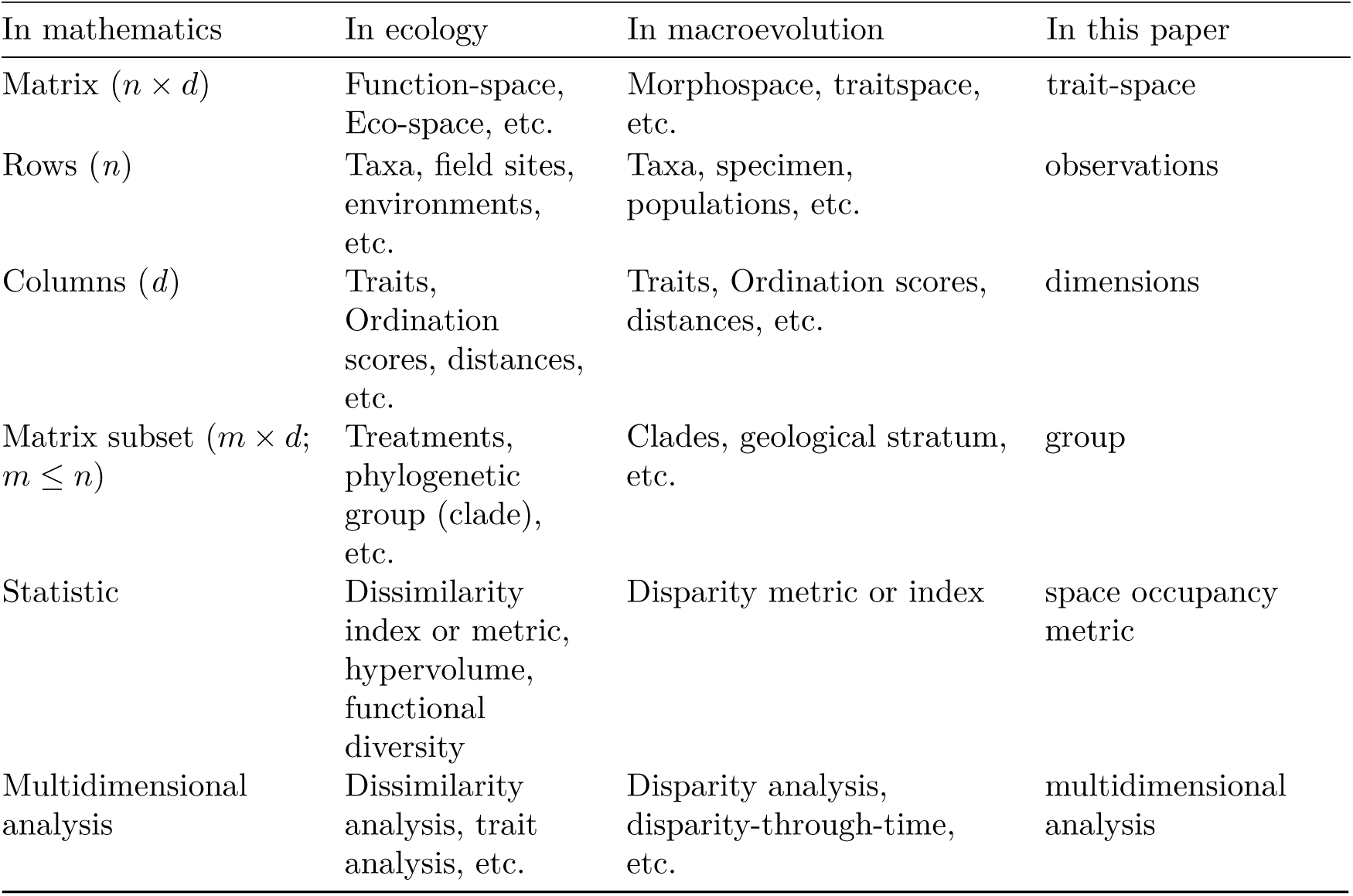
terms and equivalence between mathematics, ecology and macroevolution.

| In mathematics | In ecology | In macroevolution | In this paper |
| --- | --- | --- | --- |
| Matrix ($n \times d$) | Function-space,<br>Eco-space, etc. | Morphospace, traitspace,<br>etc. | trait-space |
| Rows ($n$) | Taxa, field sites, environments, etc. | Taxa, specimen, populations, etc. | observations |
| Columns ($d$) | Traits, Ordination scores, distances, etc. | Traits, Ordination scores, distances, etc. | dimensions |
| Matrix subset ($m \times d$; $m \leq n$) | Treatments, phylogenetic group (clade), etc. | Clades, geological stratum, etc. | group |
| Statistic | Dissimilarity index or metric, hypervolume, functional diversity | Disparity metric or index | space occupancy metric |
| Multidimensional analysis | Dissimilarity analysis, trait analysis, etc. | Disparity analysis, disparity-through-time, etc. | multidimensional analysis |

Ecologists and evolutionary biologists also often use trait-spaces with respect to the same fundamental questions: are groups overlapping in the trait-space? Are some regions of the trait-space not occupied? How do specific factors influence the occupancy of the trait-space? Studying the occupancy of trait-spaces can be achieved using disparity metrics in macroevolution (Wills 2001; Hopkins and Gerber 2017; Guillerme 2018) or comparing hypervolumes in ecology (Donohue et al. 2013; Díaz et al. 2016; Blonder 2018; Mammola 2019). However, although these space occupancy metrics are common in ecology and evolution, surprisingly little work has been published on their behaviour (but see Ciampaglio et al. 2001; Villéger et al. 2008; Mammola 2019).

Different space occupancy metrics capture different aspects of the trait-space (ciampaglio2001; Villéger et al. 2008; Mammola 2019). It may be widely-known by many in the field, but to our knowledge this is infrequently mentioned in peer-reviewed papers. First, space occupancy metrics are often named as the biological aspect they are describing (e.g. “disparity” or “functional diversity”) rather than what they are measuring (e.g. the average pairwise distances) which obscures the differences or similarities between studies. Second, in many studies in ecology and evolution, authors have focused on measuring the volume of the trait-space with different metrics (e.g. ellipsoid volume Donohue et al. 2013; hypervolume Díaz et al. 2016; Procrustes variance Marcy et al. 2016; product of variance Wright 2017). However, volume only represents a single aspects of space occupancy, disregarding others such as the density (Harmon et al. 2008) or position (Wills 2001; Ciampaglio et al. 2001). For example, if two groups occupy the same volume in trait-space, this will lead to supporting certain biological conclusions. Yet, an alternative aspect of space occupancy may indicate that the groups’ position are different; this would likely lead to a different biological conclusion (e.g. the groups are equally diverse but occupy different niches). Using metrics that only measure one aspect of the multidimensional trait-space may restrain the potential of multidimensional analysis (Villéger et al. 2008).

Here we propose a broad classification of space occupancy metrics as used across ecology and evolution and analyse their power to detect changes in trait-space occupancy in simulated and empirical data. We provide an assessment of each broad type of space occupancy metrics along with a unified terminology to foster communication between ecology and evolution. Unsurprisingly, we found no one metric describes all changes through a trait-space and the results from each metric are dependent on the characteristics of the space and the hypotheses. Furthermore, because there can potentially be an infinite number of metrics, it would be impossible to propose clear generalities to space occupancy metrics behavior. Therefore, we propose `moms`, a user-friendly tool allowing researchers to design, experiment and visualise their own space occupancy metric tailored for their specific project and helping them understanding the “null” behavior of their metrics of interest.

### Space occupancy metrics

In this paper, we define trait-spaces as any matrix where rows are observations and columns are traits. These traits can widely vary in number and types: they could be coded as discrete (e.g. presence or absence of a bone; Beck and Lee 2014; Wright 2017), continuous measurements (e.g. leaf area; Díaz et al. 2016) or more sophisticated measures (Fourier ellipses; Bonhomme et al. 2014; e.g. landmark position; Marcy et al. 2016). Traits can also be measured by using relative observations (e.g. community compositions; Jones et al. 2015) or distance between observations (e.g. Close et al. 2015). However, regardless of the methodology used to build a trait-space, three broad occupancy metrics can be measured: the volume which will approximate the amount of space occupied, the density which will approximate the distribution in space and the position which will approximate the location in space (Fig. 1; Villéger et al. 2008). Of course any combination of these three aspects is always possible.

**Figure 1:**
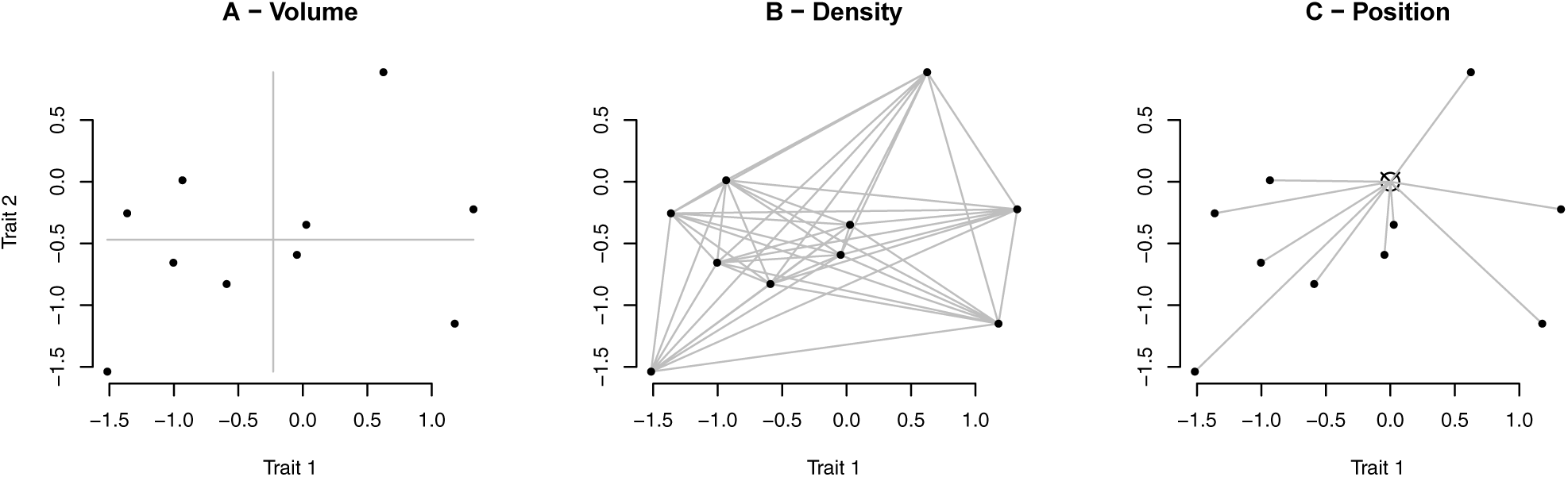
different type of information captured by space occupancy metrics. A - Volume (e.g. sum of ranges); B - Density (e.g. average squared pairwise distances); C - Position (e.g. median distance from centroid).

#### 1. Volume

Volume metrics measure the spread of a group in the trait-space. They can be interpreted as the amount of the trait-space that is occupied by observations. Typically, larger values for such metrics indicate the presence of more extreme trait combinations. For example, if group A has a bigger volume than group B, the observations in A achieve more extreme trait combinations than in B. This type of metric is widely used in both ecology (e.g. the hypervolume; Blonder 2018) and in evolution (e.g. the sum or product of ranges or variances; Wills 2001).

Although volume metrics are a suitable indicator for comparing a group’s trait-space occupancy, it is limited to comparing the range of trait-combinations between groups. Volume metrics do not take into account the distribution of the observations within a group. In other words, they can make it difficult to determine whether all the observations are on the edge of the volume or whether the volume is simply driven by small number of extreme observations.

#### 2. Density

Density metrics measure the distribution of a group in the trait-space. They can be interpreted as the distribution of the observations *within* a group in the trait-space. Groups with higher density have observations within it that tend to be more similar to each other. For example, if group A has a greater volume than group B but both have the same density, similar mechanisms could be driving both groups’ trait-space occupancy. However, this might suggest that A is older and had more time to achieve more extreme trait combinations under essentially the same process (Endler et al. 2005). Density is less commonly measured compared to volume, but it is still used in both ecology (e.g. the minimum spanning tree length; Oksanen et al. 2007) and evolution (e.g. the average pairwise distance; Harmon et al. 2008).

#### 3. Position

Position metrics measure where a group lies in trait-space. They can be interpreted as where a group lies in the trait-space either relative to the space itself or relative to another group. For example, if group A has a different position than group B, observations in A will have a different trait-combination than B.

Position metrics may be harder to interpret in multidimensional spaces (i.e. beyond left/right, up/down and front/back). However, when thinking about unidimensional data (one trait), this metric is obvious: two groups A or B could have the same variance (i.e. “volume” or spread) with the same number of observations (i.e. density) but could have a different mean and thus be in different positions. These metrics have been used in ecology to compare the position of two groups relatively to each other (Mammola 2019).

### No metric to rule them all: benefits of considering multiple metrics

The use of multiple metrics to assess trait-space occupancy has the benefit of providing a more detailed characterisation of occupancy changes. If the question of interest is, say, to look at how space occupancy changes in response to mass extinction, using a single space occupancy metric can miss part of the picture: a change in volume could be decoupled from a change in position or density in trait-space. For example, the Cretaceous-Palaeogene extinction (66 million years ago - Mya) has been linked to an increase in volume of the mammalian trait-space (adaptive radiation; Halliday and Goswami 2016) but more specific questions can be answered by looking at other aspects of trait-space occupancy: does the radiation expands on previously existing morphologies (elaboration, increase in density; Endler et al. 2005) or does it explore new regions of the trait-space (innovation, change in position; Endler et al. 2005)? Similarly, in ecology, if two groups occupy the same volume in the trait-space, it can be interesting to look at differences in density within these two groups: different selection pressure can lead to different density within equal volume groups.

Here, we provide the first interdisciplinary review of 25 space occupancy metrics that uses the broad classification of metrics into volume, density and position to capture pattern changes in trait-space. We assess the behaviour of metrics using simulations and six interdisciplinary empirical datasets covering a wide range of potential data types and biological questions. We also introduce a tool for measuring occupancy in multidimensional space (`moms`), which is a user-friendly, open-source, graphical interface to allow the tailored testing of metric behaviour for any use case. `moms` will allow workers to comprehensively assess the properties of their trait-space and the metrics associated with their specific biological question.

## Methods

We tested how 25 different space occupancy metrics relate to each other, are affected by modifications of traits space and affect group comparisons in empirical data. To do so, we performed the following steps (explained in more detail below):

1. We simulated 13 different spaces with different sets of parameters;
2. We transformed these spaces by removing 50% of the observations following four different scenarios corresponding to different empirical scenarios: randomly, by limit (e.g. expansion or reduction of niches), by density (e.g. different degrees of competition within a guild) and by position (e.g. ecological niche shift).
3. We measured occupancy on the resulting transformed spaces using eight different space occupancy metrics;
4. We applied the same space occupancy metrics to six empirical datasets (covering a range of disciplines and a range of dataset properties).

Note that the paper contains the results for only eight metrics, the results for the additional 17 metrics is available in the supplementary material 4.

### Generating spaces

We generated trait-spaces using the following combinations of size, distributions, variance and correlation:

The differences in trait-space sizes (200 × 3, 200 × 15, 200 × 50 or 200 × 150) reflects the range of dimensions in literature: “low-dimension” spaces (< 15) are common in ecology (Mammola 2019) whereas high dimension spaces ((> 100) are common in macroevolution (Hopkins and Gerber 2017). We used a range of distributions (uniform, normal or random) to test the effect of observation distributions on the metrics. We used different levels of variance for each dimensions in the spaces by making the variance on each dimension either equal (*σ*_*D*1_≃ *σ*_*D*2_≃ *σ*_*Di*_) or decreasing (*σ*_*D*1_ < *σ*_*D*2_ < *σ*_*Di*_) with the decreasing factor being either multiplicative (using the cumulative product of the inverse of the number of dimensions: 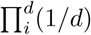 or additive 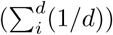. Both multiplicative and cumulative reductions of variance are used to illustrate the properties of ordinations where the variance decreases per dimensions (healy2019; and in a normal way in Multidimensional Scaling - MDS, PCO or PCoA; e.g. Close et al. 2015; in a lognormal way in principal components analysis - PCA; e.g. Marcy et al. 2016; Wright 2017). Finally, we added a correlation parameter to take into account the potential correlation between different traits. We repeated the simulation of each trait-space 20 times (resulting in 260 trait-spaces).

### Spatial occupancy metrics

We then measured eight different metrics on the resulting transformed spaces, including a new metric we produced, the average displacement, which we expect to be influenced by changes in trait-space position.

### Metric comparisons

We compared the space occupancy metrics correlations across all simulations between each pair of metrics to assess they captured signal (Villéger et al. 2008; Laliberté and Legendre 2010). We used the metrics on the full 13 trait-spaces described above. We then scaled the results and measured the pairwise Pearson correlation to test whether metrics were capturing a similar signal (high positive correlation), a different signal (correlation close to 0) or an opposite signal (high negative correlations) using the psych package (Revelle 2018).

### Changing space

To measure how the metrics responded to changes within trait-spaces, we removed 50% of observations each time using the following algorithms:

- **Randomly:** by randomly removing 50% of observations (Fig. 2-A). This reflects a “null” biological model of changes in trait-space: the case when observations are removed regardless of their intrinsic characteristics. For example, if diversity is reduced by 50% but the trait-space volume remains the same, there is a decoupling between diversity and space occupancy (Ruta et al. 2013). Our selected metrics are expected to not be affected by this change.
- **Limit:** by removing all observations with a distance from the centre of the trait-space lower or greater than a radius *ρ* (where *ρ* is chosen such that 50% observations are selected) generating two limit removals: *maximum* and *minimum* (respectively in orange and blue; Fig. 2-B). This can reflect a strict selection model where observations with trait values below or above a threshold are removed leading to an expansion or a contraction of the trait-space. Volume metrics are expected to be most affected by this change.
- **Density:** by removing any pairs of point with a distance *D* from each other where (where *D* is chosen such that 50% observations are selected) generating two density removals: *high* and *low* (respectively in orange and blue; Fig. 2-C). This can reflect changes within groups in the trait-space due to ecological factors (e.g. competition can generate niche repulsion - lower density; Grant and Grant 2006). Density metrics are expected to be most affected by this change.
- **Position:** by removing points similarly as for **Limit** but using the distance from the furthest point from the centre generating two position removals: *positive* and *negative* (respectively in orange and blue; Fig. 2-D). This can reflect global changes in trait-space due, for example, to an entire group remaining diverse but occupying a different niche. Position metrics are expected to be most affected by this change.

The algorithm to select *ρ* or *D* is described in greater detail in in the Supplementary material 1. To measure the effect of space reduction, distribution and dimensionality on the metric, we scaled the metric to be relative to the non-reduced space for each dimension distribution or number of dimensions. We subtracted the observed occupancy with no space reduction to all the occupancy measurements of the reduced spaces and then divided it by the resulting maximum observed occupancy. Our occupancy metrics where scaled between −1 and 1 with a value of 0 indicating no effect of the space reduction and *>* 0 and *<* 0 respectively indicating an increase or decrease in the occupancy metric value. We then measured the probability of overlap of the between the non-random removals (limit, density and position) and the random removals using the Bhattacharrya Coefficient (probability of overlap between two distributions; Bhattacharyya 1943).

**Figure 2:**
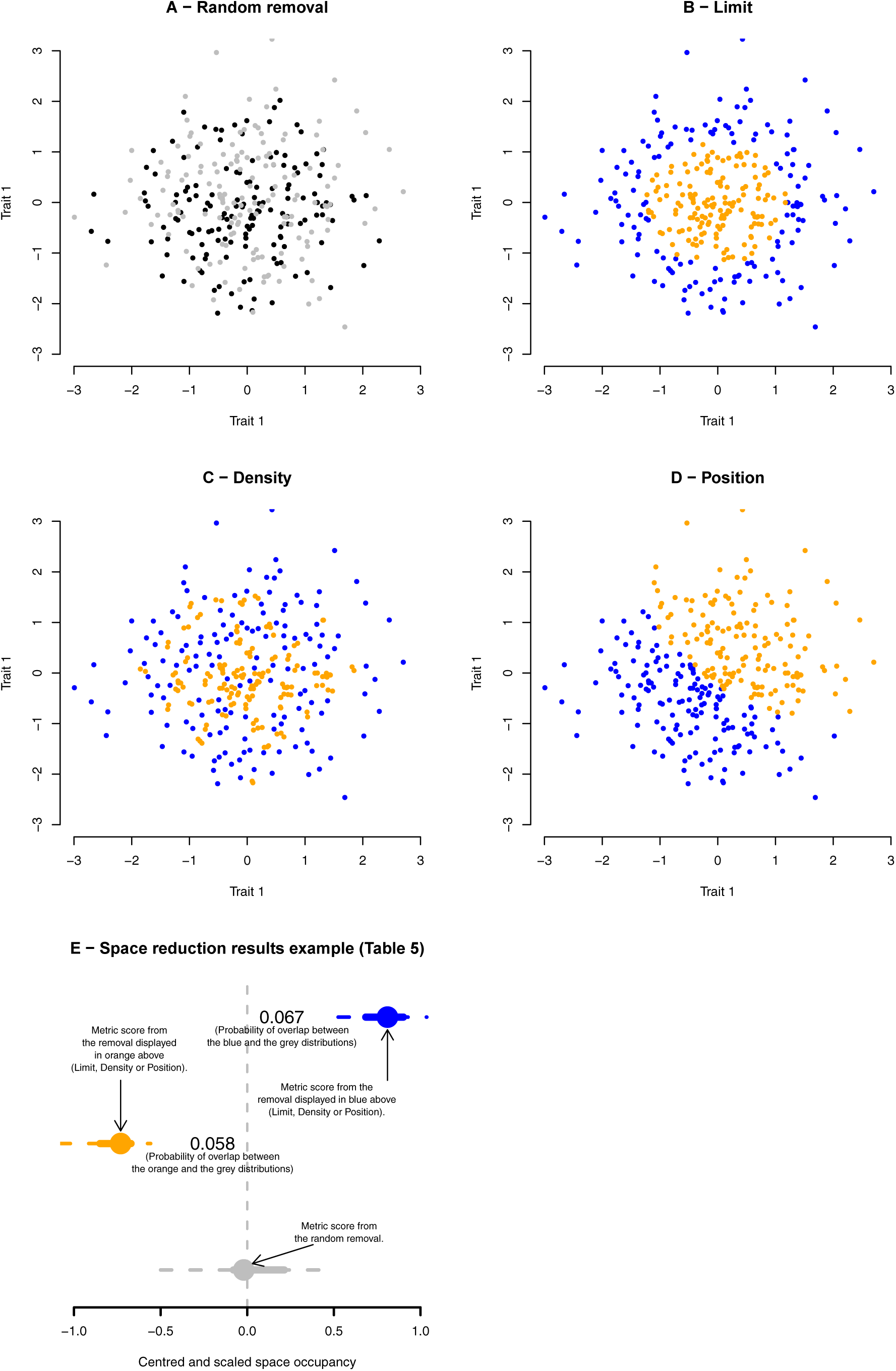
different type of space reduction. Each panel displays two groups of 50% of the data points each. Each group (orange and blue) are generated using the following algorithm: A - randomly; B - by limit (maximum and minimum limit); C - by density (high and low); and D - by position (positive and negative). Panel E represents a typical display of the reduction results displayed in Table 5: the dots represent the median space occupancy values across all simulations for each scenario of trait-space change (Table 2), the solid and dashed line respectively the 50% and 95% confidence intervals. Results in grey are the random 50% reduction (panel A). Results in blue and orange represent the opposite scenarios from panels B, C, and D. The displayed value is the probability of overlap (Bhattacharrya Coefficient) between the blue or orange distributions and the grey one.

### Measuring the effect of space and dimensionality on shifting spaces

Distribution differences and the number of dimensions can have an effect on the metric results. For example, in a normally distributed space, a decrease in density can often lead to an increase in volume. This is not necessarily true in log-normal spaces or in uniform spaces for certain metrics. Furthermore, high dimensional spaces (>10) are subject to the “curse of multidimensionality” (Chávez et al. 2001): data becomes sparser with increasing number of dimensions, such that the probability of two points A and B overlapping in *n* dimensions is the product of the probability of the two points overlapping on each dimensions 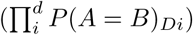. This probability decreases as a product of the number of dimensions. Therefore, the “curse” can make the interpretation of high dimensional data counter-intuitive. For example if a group expands in multiple dimensions (i.e. increase in volume), the actual hypervolume can decrease (Fig. 3 and Tables 6, 7).

**Table 2:**
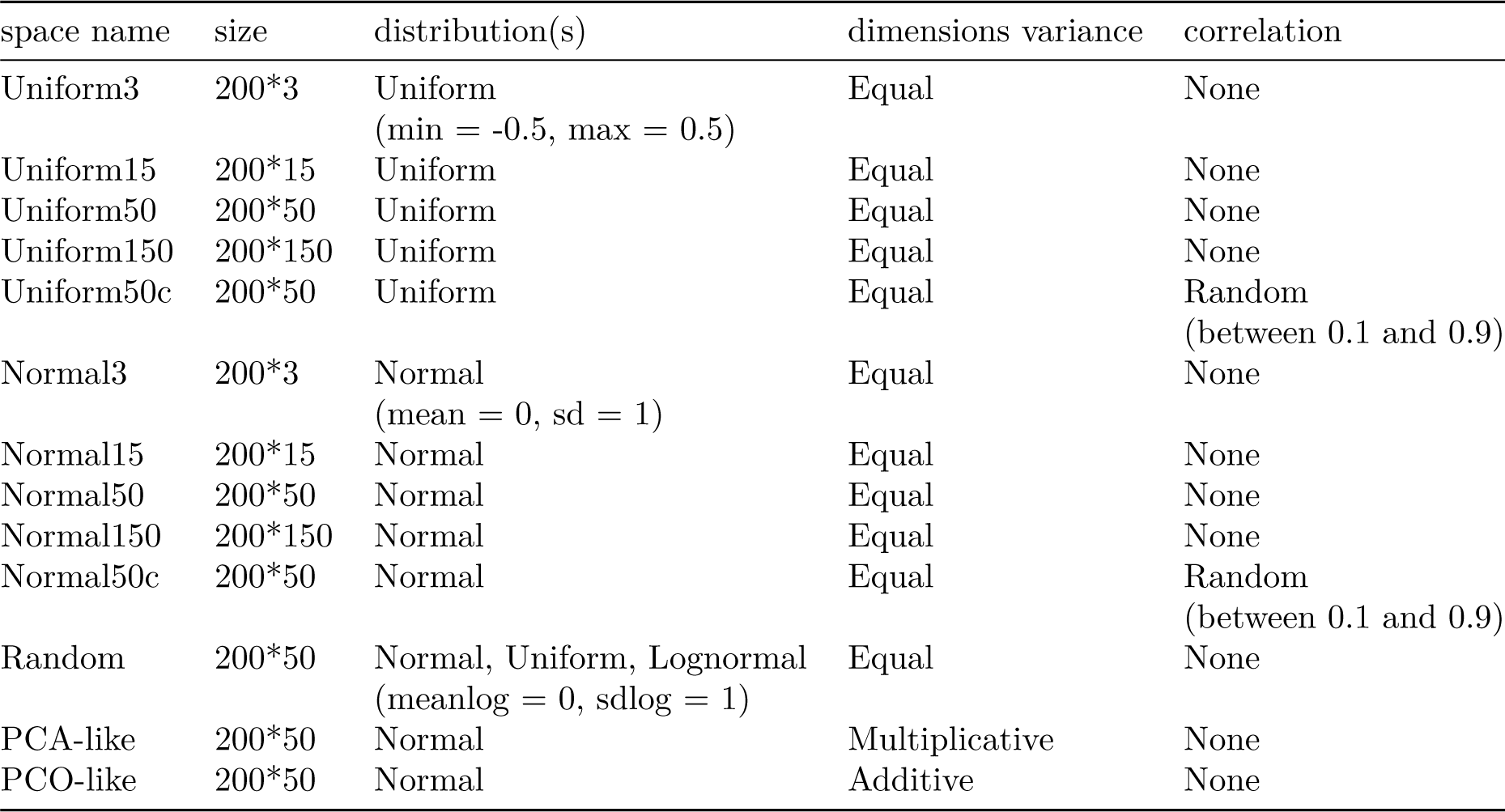
different simulated space distributions.

**Table 3:** List of metrics with *n* being the number of observations, *d* the total number of dimensions, *k* any specific row in the matrix, *Centroid* being their mean and *σ*^2^ their variance. Γ is the Gamma distribution and *λ*_*i*_ the eigen value of each dimension and *q*_*i*_ and *p*_*i*_ are any pairs of coordinates.

| Name | Definition | Captures | Source | Notes |
| --- | --- | --- | --- | --- |
| Average distance from centroid | $\frac{\sqrt{\sum_i^n (k_i - \text{Centroid}_k)^2}}{d}$ | Volume | Laliberté and Legendre (2010) | equivalent to the functional dispersion (FDis) in Laliberté and Legendre (2010) (without abundance) |
| Sum of variances | $\sum_i^d \sigma^2 k_i$ | Volume | Wills (2001) | common metric used in palaeobiology (Ciampaglio et al. 2001) |
| Sum of ranges | $\sum_i^d \ \max(d_i) - \min(d_i)\ $ | Volume | Wills (2001) | more sensitive to outliers than the sum of variances |
| Ellipsoid volume | $\frac{\pi^{d/2}}{\Gamma(\frac{d}{2}+1)} \prod_i^d (\lambda_i^{0.5})$ | Volume | Donohue et al. (2013) | less sensitive to outliers than the convex hull hypervolume (Díaz et al. 2016; Blonder 2018) |
| Minimum spanning tree average distance | $\frac{\sum (\text{branch length})}{n}$ | Density | Oksanen et al. (2007) | similar to the unscaled functional evenness (Villéger et al. 2008) |
| Minimum spanning tree distances evenness | $\frac{\sum \min \left( \frac{\text{branch length}}{\sum \text{branch length}} \right) - \frac{1}{n-1}}{1 - \frac{1}{n-1}}$ | Density | Villéger et al. (2008) | the functional evenness without weighted abundance (FEve; Villéger et al. 2008) |
| Average nearest neighbour distance | $\min \left( \sqrt{\sum_i^n (q_i - p_i)^2} \right) \times \frac{1}{n}$ | Density | Foote (1990) | the density of pairs of observations in the trait-space |
| Average displacement | $\frac{\sqrt{\sum_i^n (k_i)^2}}{\sqrt{\sum_i^n (k_i - \text{Centroid}_k)^2}}$ | Position | This paper | the ratio between the observations' position from their centroid and the centre of the trait-space. A value of 1 indicates that the observations' centroid is the centre of the trait-space |

**Table 4:**
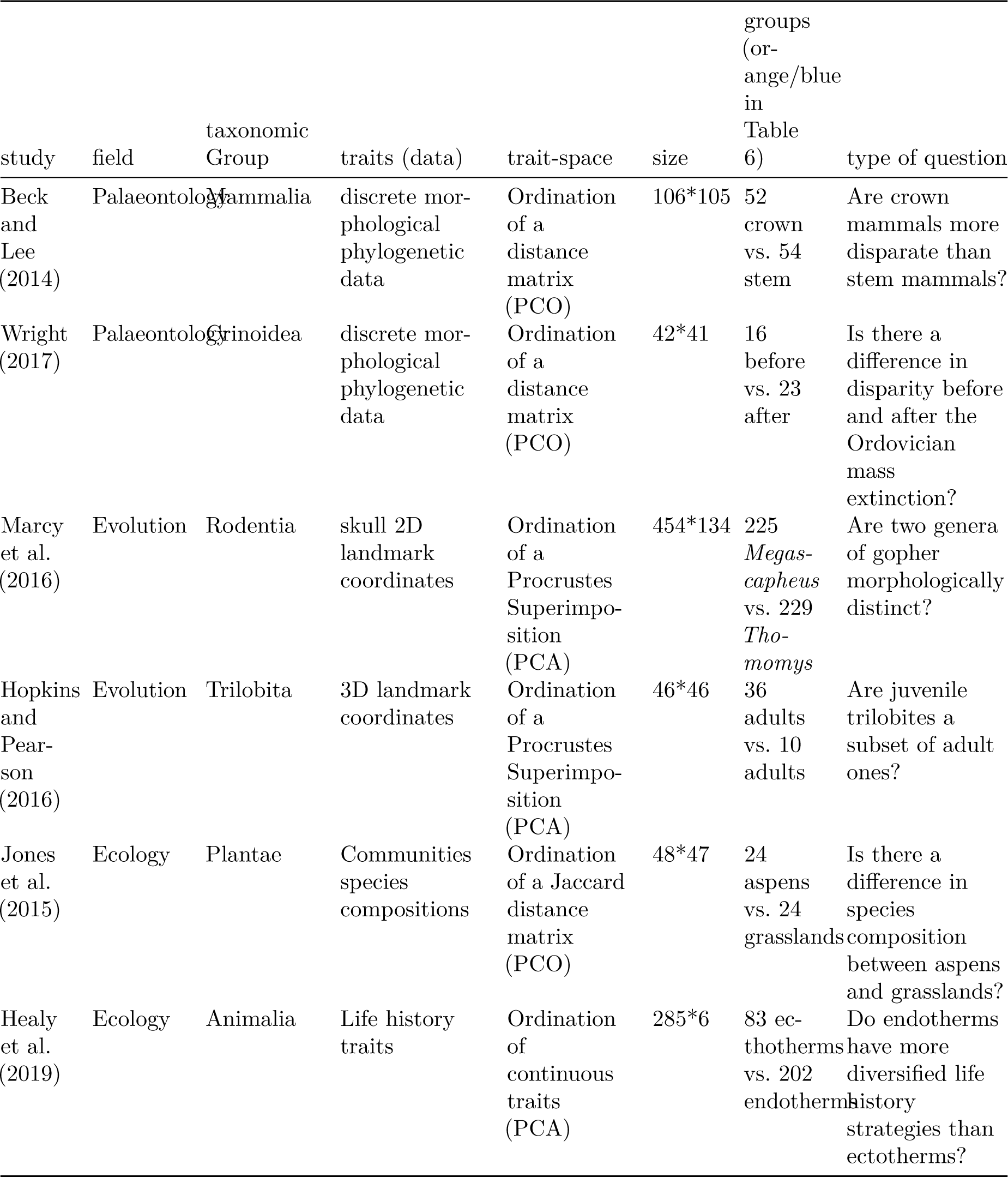
details of the six empirical trait-spaces.

**Table 5:**
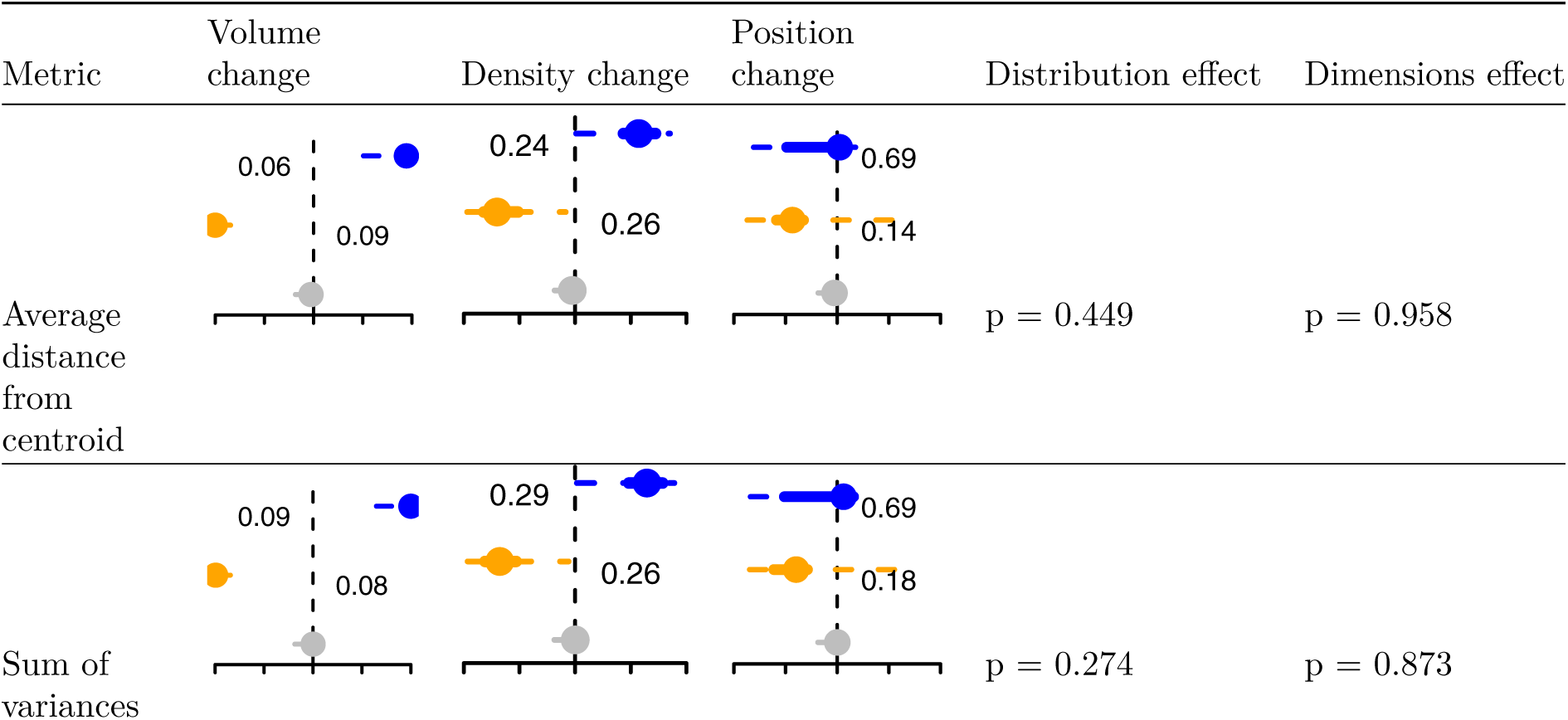
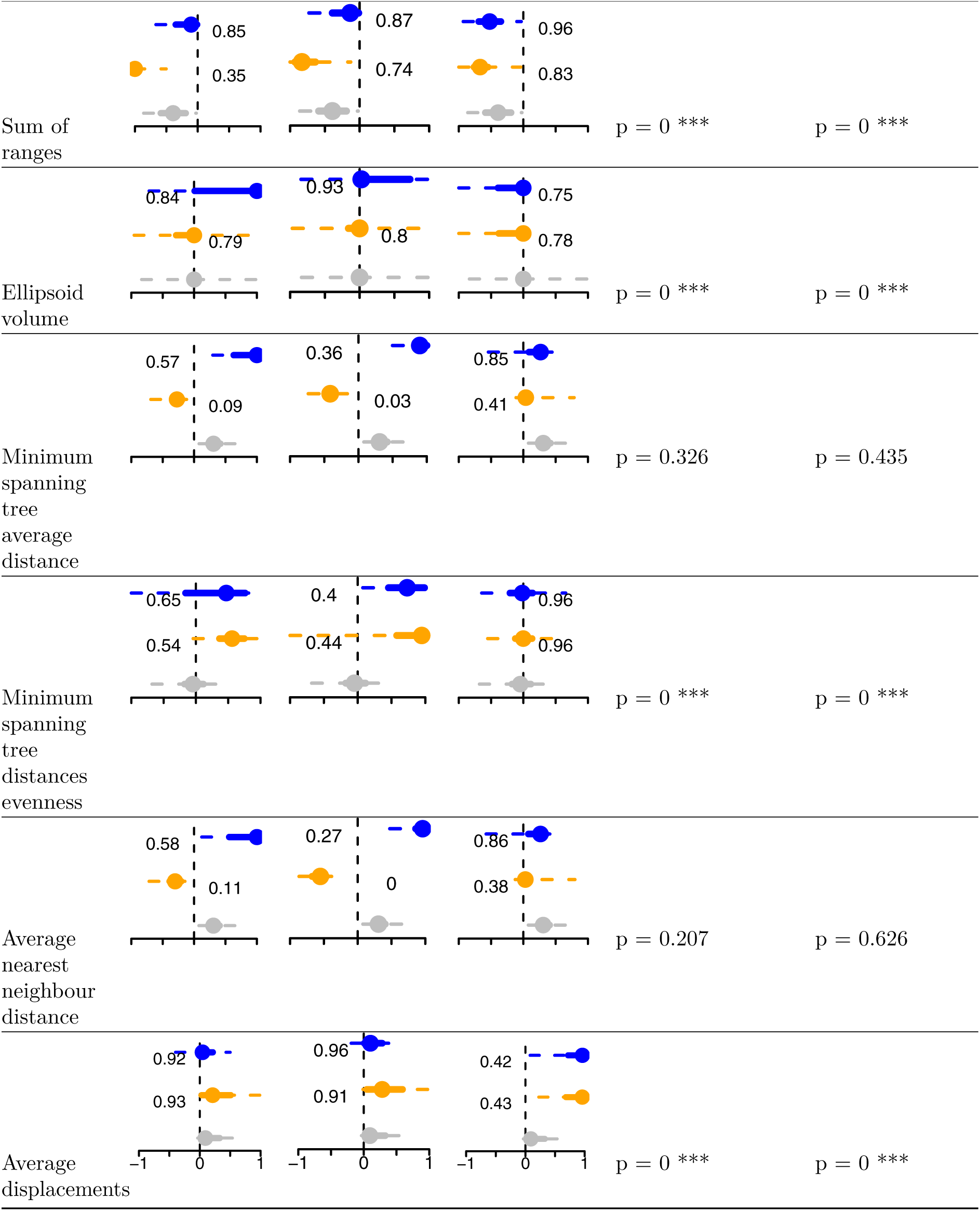
Results of the effect of space reduction, space dimension distributions and dimensions number of the different space occupancy metrics. See Fig. 2 for interpretation of the figures. *p*-values for distribution effect and dimensions effect represents respectively the effect of the ANOVAs space occupancy ∼ distributions and space occupancy ∼ dimensions (0 ‘***’ 0.001 ‘**’ 0.01 ‘*’ 0.05 ‘.’ 0.1” 1).

**Table 6:**
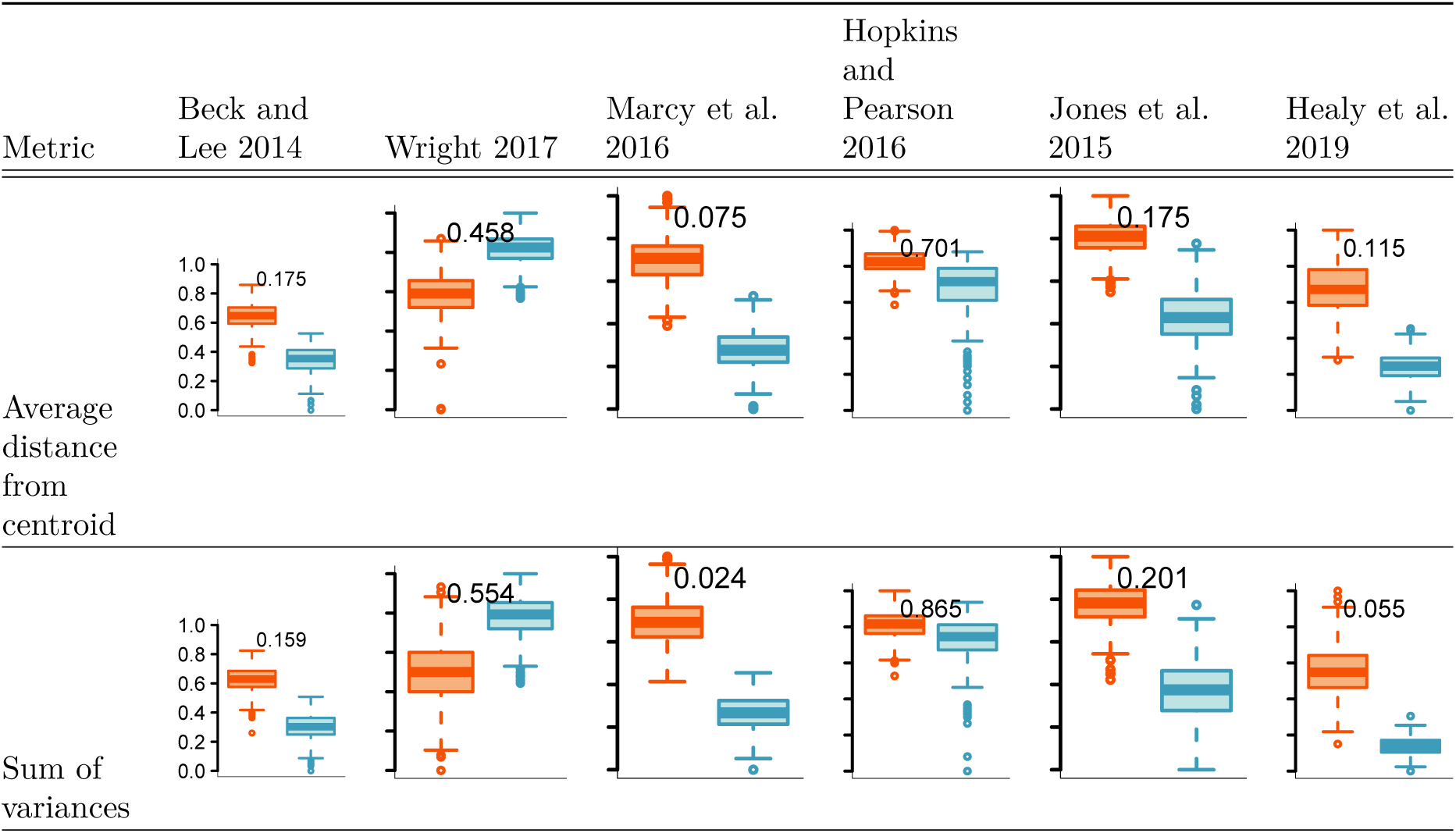
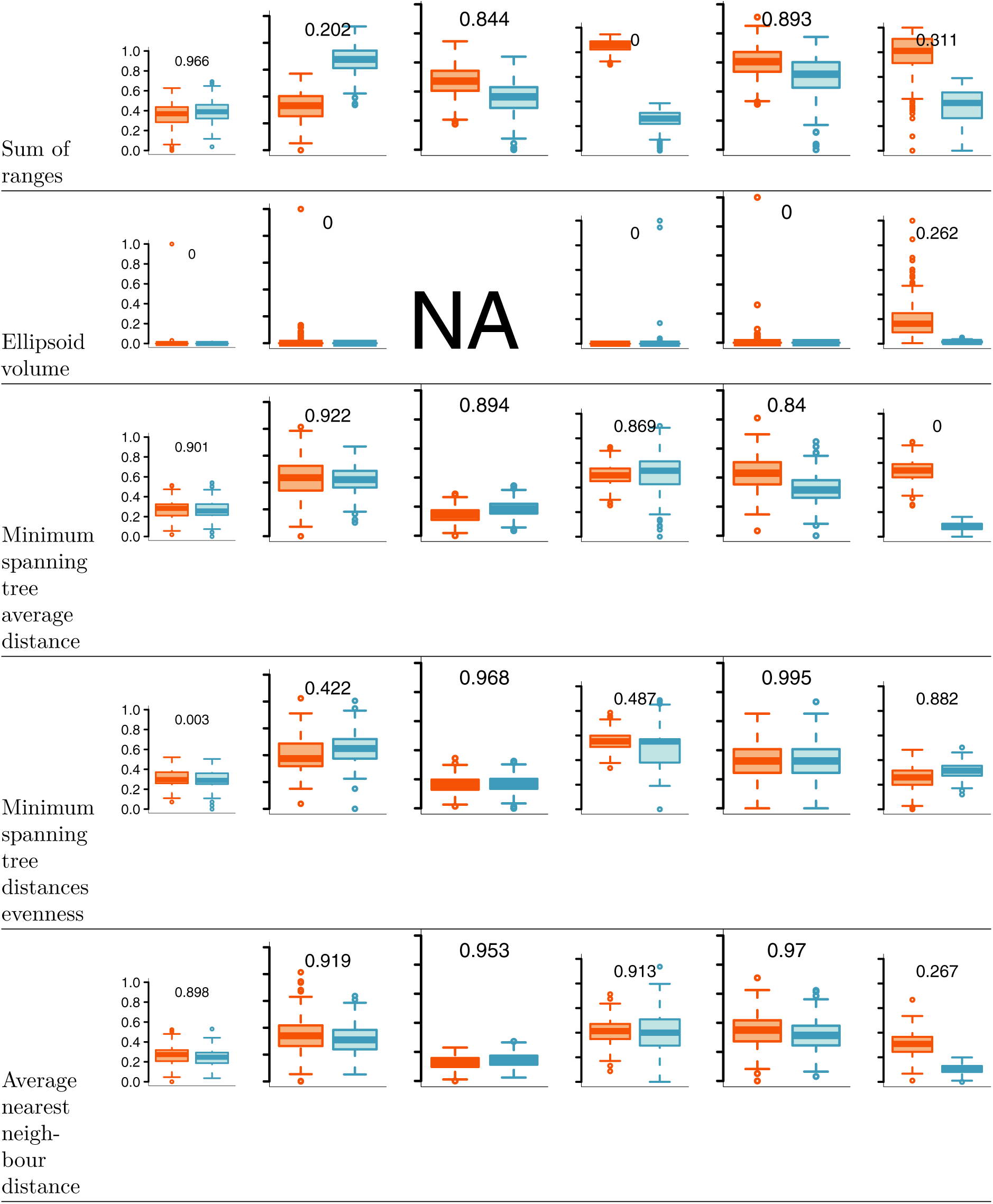
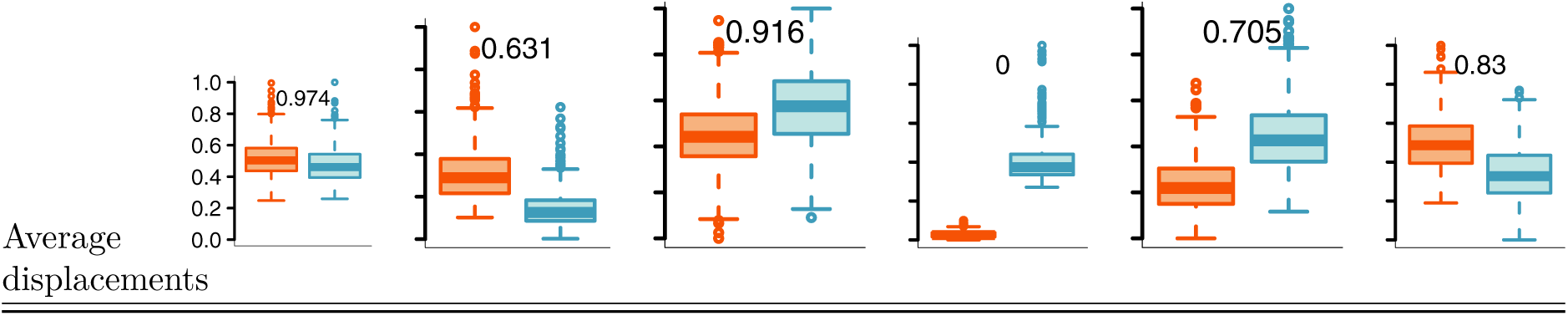
Comparisons of pairs of groups in different empirical trait-spaces. NAs are used for cases where space occupancy could not be measured due to the curse of multidimensionality. The displayed values are the probability of overlap between both groups (Bhattacharrya Coefficient).

**Figure 3:**
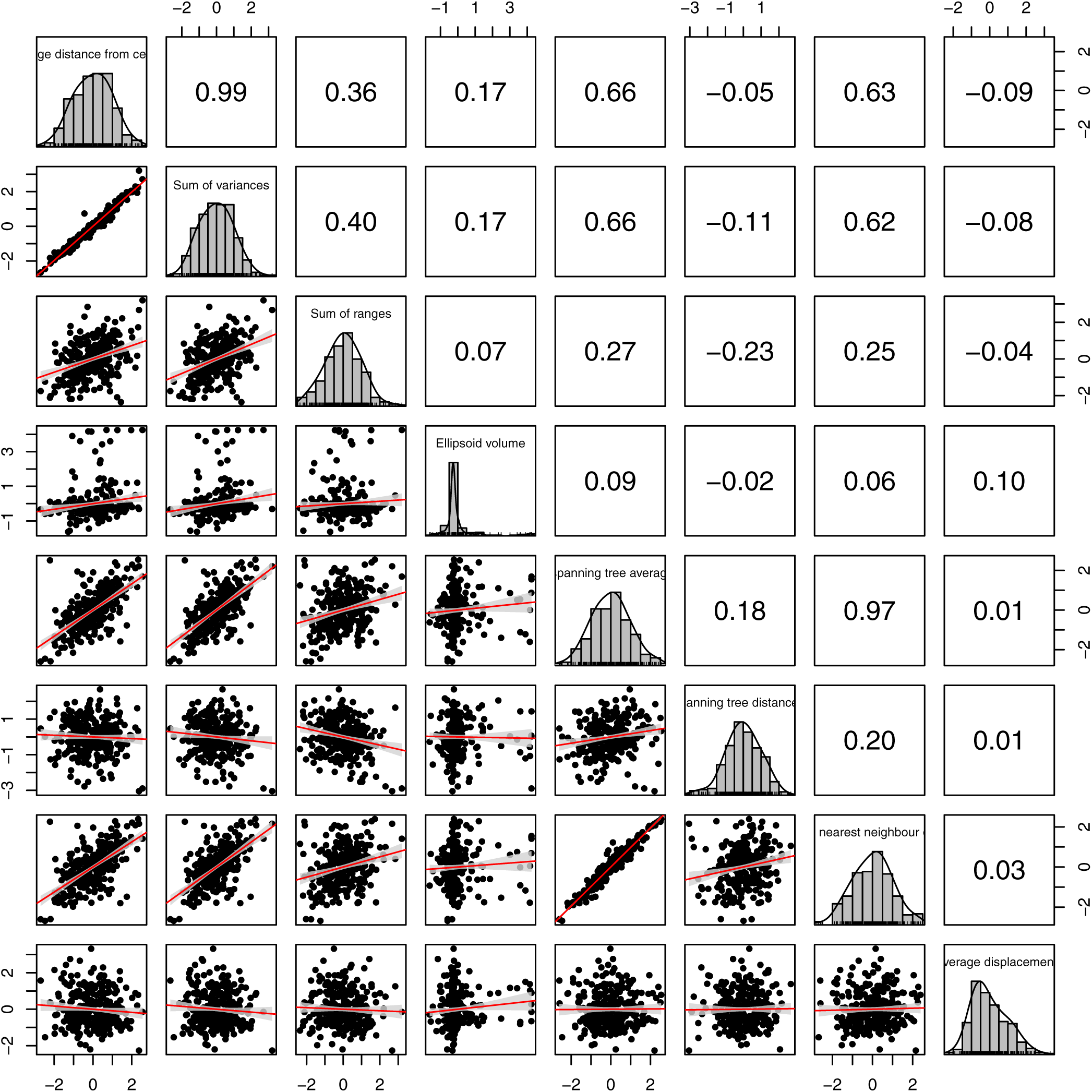
pairwise correlation between the scaled metrics. Numbers on the upper right corner are the Pearson correlations. The red line are linear regressions (with the confidence intervals in grey).

We measured the effect of space distribution and dimensionality using an ANOVA (*occupancy ∼ distribution* and *occupancy ∼ dimensions*) by using all spaces with 50 dimensions and the uniform and normal spaces with equal variance and no correlation with 3, 15, 50, 100 and 150 dimensions (Table 2) for testing respectively the effect of distribution and dimensions. The results of the ANOVAs (*p*-values) are reported in Table 5 (see supplementary material 1 for the full ANOVA result tables).

### Empirical examples

We analysed the effect of the different space occupancy metrics on six different empirical studies covering a broad range of fields that employ trait-space analyses (palaeobiology, macroevolution, evo-devo, ecology, etc.). For each of these studies we generated trait-spaces from the data published with the papers. We divided each trait-spaces into two biologically-relevant groups and tested whether the metrics differentiated the groups in different ways. Both the grouping and the questions where based on a simplified version of the topics of these papers (with no intention to re-analyse the data but to be representative of the diversity of questions in ecology and evolution). The procedures to generate the data and the groups varies from one study to the other but is detailed and fully reproducible in the supplementary materials 2.

For each empirical trait-space we bootstrapped each group 500 times (Guillerme 2018) and applied the eight space occupancy metric to each pairs of groups. We then compared the means of each groups using the Bhattacharrya Coefficient (Bhattacharyya 1943).

## Results

### Metric comparisons

All the metrics were either positively correlated (Pearson correlation of 0.99 for the average distance from centroid and sum of variance or 0.97 for the average nearest neighbour distance and minimum spanning tree average length; Fig. 3) or somewhat correlated (ranging from 0.66 for the sum of variances and the ellipsoid volume to −0.09 between the average displacement and the average distance from centroid; Fig. 3). All metrics but the ellipsoid volume were normally (or nearly normally) distributed (Fig. 3). More comparisons between metrics are available in the supplementary materials 3.

### Space shifting

As expected, some different metrics capture different aspects of space occupancy. However, it can be hard to predict the behaviour of each metric when 50% of the observations are removed. We observe a clear decrease in median metric in less than a third of the space reductions (10/36).

In terms of change in volume, only the average distance from centroid and the sum of variances seem to capture a clear change in both directions. However, the increase in volume does not correspond to an *actual* increase in volume in the trait-space (i.e. the volume from the blue observations in Fig. 2-B is equivalent to the one in Fig. 2-A). In terms of change in density, only the minimum spanning tree average distance and the average nearest neighbour distance seem to capture a clear change in both directions. In terms of change in position, only the average displacement metric seems to capture a change a clear change in direction (albeit not in both directions). This is not surprising, since the notion of positions becomes more and more complex to appreciate as dimensionality increases (i.e. beyond left/right, up/down and front/back).

### Empirical example

Similarly as for the simulated results, the empirical ones indicate that there is no perfect one-size-fit all metric. For all eight metrics (expect the ellipsoid volume) we see either one group or the other having a bigger mean than the other and no consistent case where a group has a bigger mean than the other for all the metrics. For example, in the Beck and Lee (2014)’s dataset, there is a clear non-overlap in space occupancy volume using the average distance from centroid or the sum of variances (overlaps of respectively 0.175 and 0.159) but no overlap when measuring the volume using the sum of ranges (0.966). However, for the Hopkins and Pearson (2016)’s dataset, this pattern is reversed (no clear differences for the average distance from centroid or the sum of variances - 0.701 and 0.865 respectively) but a clear difference for the sum of ranges (0). Furthermore, for each dataset, the absolute differences between each groups is not consistent depending on the metrics. For example, in Hopkins and Pearson (2016)’s dataset, the orange group’s mean is clearly higher than the blue one when measuring the sum of ranges (0) and the inverse is true when measuring the average displacement (0).

## Discussion

Here we tested 25 metrics of trait-space occupancy on simulated and empirical datasets to assess how each metric captures changes in trait-space volume, density and position. Our results show that the correlation between metrics can vary both within and between metric categories (Fig. 3), highlighting the importance of understanding the metric classification for the interpretation of results. Furthermore, our simulations show that different metrics capture different types of trait-space change (Table 5), meaning that the use of multiple metrics is important for comprehensive interpretation of trait-space occupancy. We also show that the choice of metric impacts the interpretation of group differences in empirical datasets (Table 6), again emphasizing that metric choice has a real impact on the interpretation of specific biological questions

### Metrics comparisons

Metrics within the same category of trait-space occupancy (volume, density or position) do not have the same level of correlation with each other. For example, the average distance from centroid (volume) is highly correlated to the sum of variances (volume - correlation of 0.99) and somewhat correlated with the minimum spanning tree average distance (density - correlation of 0.66) but poorly with the ellipsoid volume (volume - correlation of 0.17) and the minimum spanning tree distances evenness (density - correlation of −0.05). Furthermore, the fact that we have such a range of correlations for normal distributions suggests that each metric can capture different summaries of space occupancy ranging from obvious differences (for metrics not strongly correlated) to subtle ones (for metrics strongly correlated).

### Space shifting

Most metrics capture no changes in space occupancy for the “null” (random) space reduction (in grey in Table 5). This is a desirable behaviour for space occupancy metrics since it will likely avoid false positive errors in empirical studies that estimate biological processes from space occupancy patterns (e.g. competition Brusatte et al. 2008, convergence Marcy et al. (2016), life history traits Healy et al. (2019)). However, the average nearest neighbour distance and the sum of ranges have a respectively positive and negative “null” median. This is not especially a bad property but it should be kept in mind that even random processes can increase or decrease these metric value.

Regarding the changes in volume, the sum of variances and the average distance from centroid are good descriptors (Table 5). However, as illustrated in the 2D examples in Fig. 2-B only the blue change results (maximum limit - Table 5) should not result in a direct change in volume since the trait-space is merely “hollowed” out. That said, “hollowing” is more hard to conceptualise in many dimensions and the metrics can still be interpreted for comparing groups (orange has a smaller volume than blue).

Regarding changes in density, the average nearest neigbhour distance and the minimum spanning tree average distance consistently detect changes in density with more precision for low density trait-spaces (in blue in Table 5). However, we can observe some degree of correlation between the changes in density and the changes in volume for most metric picking either signal. This could be due to the use of normally distributed spaces where a change in density often leads to a change in volume. This is not necessary the case with empirical data.

Regarding the changes in position of the trait-space, all but the average displacement metric seems to not be able to distinguish between a random change and a displacement of the trait-space (Table 5). Furthermore, the average displacement metric does not distinguish between and positive or a negative displacement of the trait-space: this might be due to the inherent complexity of *position* in a multidimensional trait-space.

### Empirical examples

Although most differences are fairly consistent within each dataset with one group having a higher space occupancy score than the other for multiple metrics, this difference can be more or less pronounced within each dataset (ranging from no to nearly full overlap - BC *∈* (0; 0.995)) and sometimes even reversed. This indicates that opposite conclusions can be drawn from a dataset depending on which space occupancy metric is considered. For example, in Wright (2017), crinoids after the Ordovician mass extinction have a higher median metric value for all metrics but for the average displacement. These differences depending on the metrics are also more pronounced in the empirical datasets where the observations per group are unequal (Hopkins and Pearson 2016; Healy et al. 2019). ### Caveats

While our simulations have been useful to illustrate the behavior of diverse space occupancy metrics, they have several caveats. First, the simulated observations in the trait-spaces are independent. This is not the case in biology where observations can be spatially (Jones et al. 2015) or phylogenetically correlated (e.g. Beck and Lee 2014). Second, the algorithm used to reduce the trait-spaces might not always accurately empirical changes. This might favour some specific metrics over others, in particular for the changes in density that modifies the nearest neighbour density rather than changing the global density. This algorithmic choice was made in order to not confound changes in density along with changes in volume. However, the results presented here probably capture the general behaviour of each metric since results are consistent between the simulated and empirical analysis. Furthermore, `moms` allows to test the caveats mentioned above by uploading empirical trait-space.

### Suggestions

We insist that no metric is better than the next one and that researchers should use the most appropriate metrics based on the metric and trait-space properties as well as their specific biological question. However, following the findings of this study we suggest several points:

First, we suggest using multiple metrics to tackle different aspects of the trait-space. This follows the same logical thinking that the mean might not be sufficient to describe a distribution (e.g. the variance might be good additional descriptor). Although using multiple metrics is not uncommon in macroevolutionary studies (e.g. Halliday and Goswami 2016) or in ecology (Mammola 2019), they often do not cover contrasted aspects of the trait-space.

Second, we suggest selecting the metrics that best help answering the biological question a hand. If one studies an adaptive radiation in a group of organisms, it is worth thinking what would be the expected null model: would the group’s volume increase (radiation in all directions), would it increase in density (niche specialisation) or would it shift in position (radiation into a new set of niches)?

Third, we suggest to not name metrics as the biological aspect they are describing (e.g. “disparity” or “functional dispersion”) but rather what they are measuring (e.g. “sum of dimensions variance”). We believe this will allow both a clearer understanding of what *is* measured and a better communication between ecology and evolution research where metrics can be similar but have different names (Fig. 3).

Multidimensional analyses have been acknowledged to be an essential tool-kit modern biology but can often be counter-intuitive (Chávez et al. 2001). It is thus crucial to accurately describe patterns in multidimensional trait-spaces to be able to link them to biological processes. When summarising trait-spaces, it is important to remember that a pattern captured by a specific space occupancy metric is often dependent on the properties of the trait-space and of the particular biological question of interest. We believe that having a clearer understanding of both the properties of the trait-space and the associated space occupancy metrics (e.g. using `moms`) as well as using novel space occupancy metrics to answer specific questions will be of great use to study biological processes in a multidimensional world.

## Acknowledgements

We thank Natalie Jones and Kevin Healy for helping with the empirical ecological datasets. We acknowledge funding from the Australian Research Council DP170103227 and FT180100634 awarded to VW.

## Authors contributions

TG, MNP, AEM and VW designed the project. TG and AEM collected the empirical dataset. TG ran the analyses and designed the software. TG, MNP, AEM and VW wrote the manuscript.

## Data Availability, repeatability and reproducibility

The raw empirical data is available from the original papers (Beck and Lee 2014; Jones et al. 2015, Marcy et al. (2016); Hopkins and Pearson 2016; Wright 2017; Healy et al. 2019). The subsets of the empirical data used in this analysis are available on figshare DOI: 10.6084/m9.figshare.9943181.v1. The modified empirical data are available in the package accompanying this manuscript (`data(moms ∷demo_data)`). This manuscript (including the figures, tables and supplementary material) is repeatable and reproducible by compiling the vignette of the GitHub `moms R` package. The code for the `moms` shiny app is available from the GitHub `moms R` package.

